# RF-SIRF defining reversed DNA replication forks with single-cell and spatio-temporal resolution reveals a replication stress specific epigenetic code

**DOI:** 10.1101/2025.04.21.649830

**Authors:** Sunetra Roy, Morgan Fimreite, Yue Chen, Francesca Citron, Giulio Draetta, Katharina Schlacher

## Abstract

DNA replication stress responses are key genomic stability guardians critical during development, aging, hematopoiesis, disease suppression and cancer therapy response. Reversed forks (RF) form at stalled DNA replication forks as a distinct four-way DNA structure to protect against formation and exposure to toxic DNA lesions. So far, prevailing methods to measure RFs involve specialized electron microscopy precluding studies within their cellular context. Here we describe an in-situ method to quantitatively measure RFs by harnessing intrinsic bio-physical properties of this distinct DNA structure (RF-SIRF). RF-SIRF reveals that RFs accumulate at the nuclear periphery and form predominantly during early-mid S-phase of the cell cycle. Importantly, RFs are chromatinized and utilize an epigenetic replication stress code distinct from transcription that explains how DNA stress response proteins are recruited to RFs. Collectively, RF-SIRF enables robust, quantitative, temporal and spatial analyses of RFs and associated proteomics, empowering advanced cellular investigations of DNA replication stress responses.

## Introduction

Accumulating evidence shows reversed replication forks (RF) play critical roles in diverse biological settings contributing to aging, hematopoiesis, cellular senescence, stemness, and cellular survival with cancer therapies. RF are formed during stress conditions as a protective measure that promotes DNA damage tolerance in actively replicating cells to avoid exposure to DNA lesions and formation of toxic DNA double strand breaks ^1–4^. Thus, RF are of particular importance in highly proliferating cells, those exposed to DNA damage and those that are under metabolic stress. RFs accumulate in both aging and embryonic stem cells ^5, 6^, are needed to suppress myeloid and lymphoid dysregulation in both long-term hematopoietic stem cells as well as progenitor cells ^6^, and form at telomers as implied in preventing replicative senescence ^7, 8^. On the flip side, RFs also can be detrimental within certain genetic backgrounds leading to telomere catastrophe as well as genome-wide instability such as in BRCA1/2 fork protection defective cancer cells, where formation of RFs controls cellular sensitivity and resistance to standard of care and PARP inhibitor cancer therapies ^9–17^.

Fork reversal involves annealing of newly replicated DNA strands, which converts a three-way into a four-way DNA junction and forms a double-stranded DNA end within the newly regressed arm. *In vitro*, positive super-coiling of DNA provides sufficient free energy to achieve fork reversal ^17^. In cells, fork reversal is achieved by designated translocases including SMARCAL1, HLTF and ZRAN3B, with added DNA remodeling proteins FHB1, FANCM helicases, RAD51 recombinase and RAD51 paralogs being also implied in the process ^18–24^. SMARCAL1 holds a biochemically distinct role in annealing RPA coated stalled leading strands and displays a somewhat dominant function under several cellular contexts including in BRCA2 defective cells and at telomers ^12–15, 20, 25–28^. The recent discovery of SMARCAL1 status controlling inflammation and its use as a biomarker for immune checkpoint therapy response corroborates this notion and further emphasizes the importance of RF reactions in diverse cancer cell therapy settings ^26^.

Despite their key importance in both health and disease, robust tools to examine RF structures in cells are limited. Surrogate RF assays that indirectly measure RF include DNA fiber analysis, 2D-gel electrophoreses and immunofluorescence approaches ^1, 2, 22^. Yet, these are limited in efficiency, sensitivity, resolution, specificity, or quantitative robustness, restricting their broad usage. To date, the prevailing accepted assay to measure RFs with sufficient resolution and robustness is by electron microscopy (EM) ^1, 2, 22, 29^. EM visualizes RF structures, provides quantitative information regarding the relative abundance of RF and allows for single-molecule size measurements. However, this *in vitro* method is technically challenging requiring sensitive processing steps, biophysics expertise and specialized instrumentation, making the assay a time-consuming and costly effort that is prohibitive for most laboratories. Moreover, it is a bulk-DNA technique that lacks single-cell resolution and spatial context about the cellular macro-environment that defines fork reversal.

Given the challenges to examining RFs, it is enabling to develop alternative, robust, sensitive and reproducible methods to assess fork reversal in cells. Here, we directly harness reversed DNA fork structure that enforces the proximity of nascently replicated leading and lagging strand to develop a robust reporter to examine fork reversal by RF-SIRF enabling the quantitative, sensitive, and reproducible analysis of reversed forks within single-cells.

## Results

### RF-SIRF signals are elevated under conditions permissive for fork reversal

We aspired to devise an efficient, robust and technically amicable assay system to detect reversed DNA replication forks by considering the bio-physical aspects of these DNA structures. In a canonical Y-form DNA replication fork, the newly synthesized strands on the leading and lagging strand templates are spatially separated (Fig. 1a, left panel). In contrast, reversed forks form a four-way DNA junction containing a regressed arm, in which the nascently replicated leading and lagging DNA strands are annealed, thus are in direct proximity to each other (Fig. 1a, right panel, green DNA strands).

**Fig. 1.**
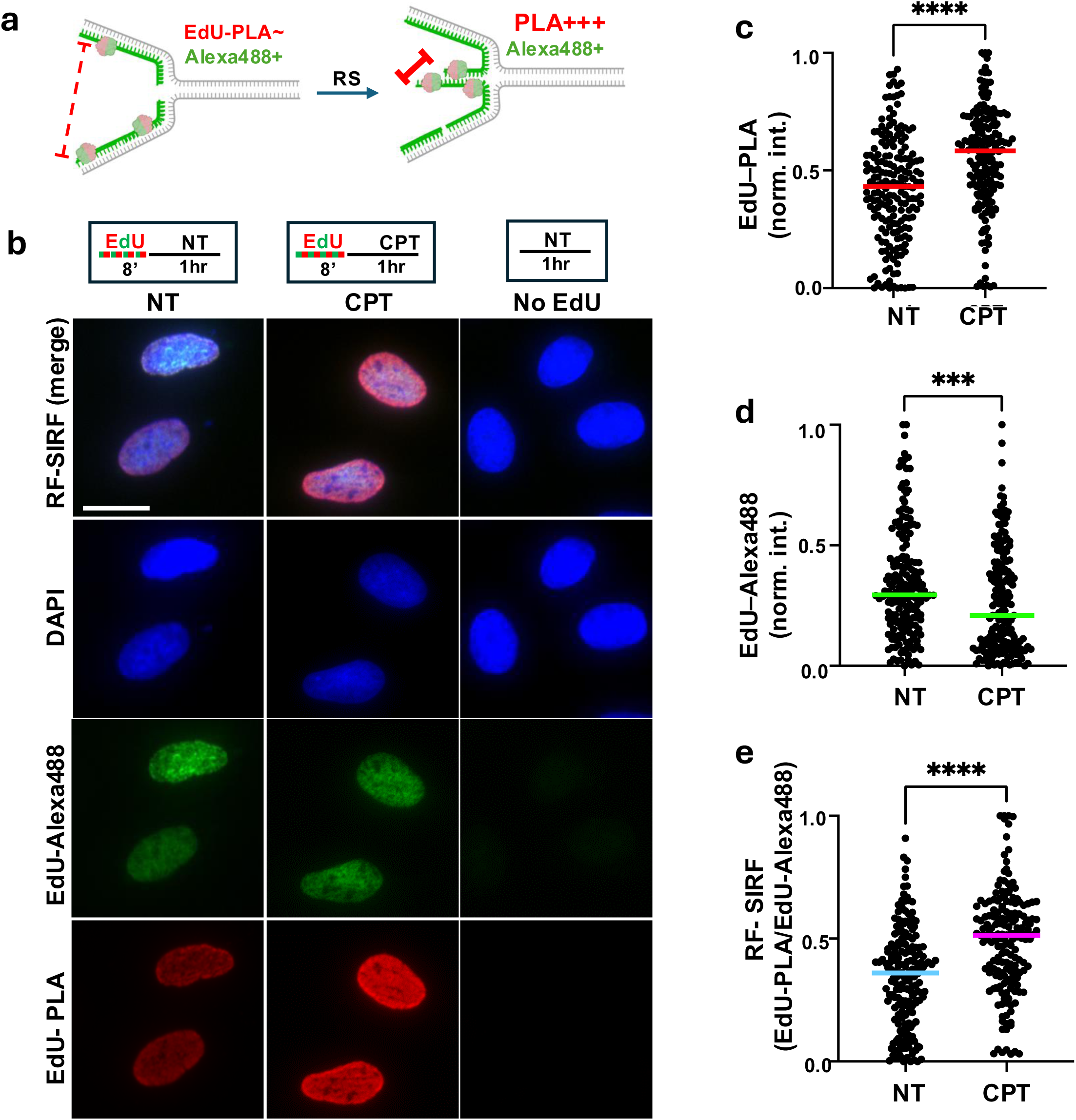
RF-SIRF signals are elevated under conditions promoting reversed forks. **a.** Schematic of principle behind RF-SIRF. Replication stress (RS) promotes fork reversal. Consequently, EdU moieties (red and green bubbles, labeled with Alexa488 and biotin) on nascent DNA (green) are in closer proximity on reversed forks where nascent leading and lagging strands (green) annealed, allowing for detection by proximity ligation assay (EdU-PLA against biotinylated EdU). EdU-Alexa488 intensities are independent of EdU-proximity and serve as substrate control. Sketch created with http://BioRender.com. **b.** Representative images of RF-SIRF. NT (no treatment); CPT (camptothecin, 50nM, 1 hour), No EdU (negative control). Top, experimental schematic. DAPI (blue) denotes nucleus. EdU-PLA (red) shows PLA against biotinylated EdU, EdU-Alexa488 (green) shows EdU with Alexa488-Click-it. Scale bar denotes 20mm. **c.** Scatter plot of EdU-PLA intensities. **d.** Scatter plot of EdU-Alexa488 intensities. **e.** Scatter plot of RF-SIRF (EdU-PLA/EdU-Alexa488). NT, without replication stalling treatment (n=190), CPT, with camptothecin promoting fork reversal (n= 178). Bars denote median of each data set. Data is derived from three independent biological repeats, *p*-values are derived using the Mann-Whitney test. \*\*\*\**p*<0.0001, \*\*\**p*<0.001

Proximity ligation assay (PLA, Extended Data Fig. 1a) exploits the spatial distance of two antibody-directed epitopes to yield a fluorescent signal only if the two epitopes are within close proximity (nominally <40nm) (Extended Data Fig. 1a) ^30–32^. Based on the unique structural elements contained within reversed fork, we reasoned that the distance between the newly incorporated Ethynyl-2’-deoxyuridine (EdU) residues is reduced in the regressed arm of the reversed fork (Fig. 1a, red-green bubble, right panel), compared to a canonical Y-shaped structure at non-stressed forks (Fig. 1a, red-green bubble, left panel). We therefore hypothesized that reversed replication forks will produce a greater EdU-PLA signal compared to Y-forks (Fig. 1a). This hypothesis builds upon and extends our previous reports on SIRF, which is an in situ assay to detect proteins at nascent replication forks using a modified PLA approach ^33–36^. For the development of SIRF, in which we biotinylate EdU *via* Click-it chemistry, we showed a direct correlation between the proximity of biotin-EdU moieties on nascently replicated DNA and the productivity of a PLA signals when using primary antibodies against biotin-EdU ^36^. Importantly, protein-SIRF signals are compared and normalized to the total amount of incorporated EdU to differentiate between signal changes caused by changes in EdU substrates (e.g. amounts of nascent DNA) from those caused by an actual loss of proximity (e.g. loss of protein-DNA interaction). To achieve this, we perform simultaneous Click-it chemistry with biotin-azide and Alexa488-azide so that both biotin and Alexa488 directly correlate to the total amount of EdU moieties present. Given that the Click-it reaction is a covalent chemical reaction that depends on substrate availability but not proximity, Alexa488 serves as a direct measure and normalization control for the total amount of incorporated EdU. In contrast, EdU-PLA signals using anti-biotin antibodies depend on the proximity between the nascently incorporated EdU moieties, which we suggest will be greater at RFs compared to ongoing Y-forks (Fig. 1a).

To test our RF hypothesis, we performed a reversed fork (RF)-SIRF whereby PLA against biotinylated EdU is performed and compared to the total amount of EdU as measured by Alexa488-co-Click-it (Fig.1a, Extended Data Fig. 1a). To devise a robust and specific assay system we considered three distinct features. First, for RF-SIRF we used cells and replication stalling conditions extensively validated to promote fork reversal ^12, 15, 22, 37, 38^. Second, EdU is incorporated within a short 8’ pulse only prior to exposure to the reversal-inducing DNA replication stress. This ensures that a change in nascent DNA signal reflects fork regression of previously incorporated EdU molecules prior to reversal. Third, EdU-Alexa488 is used to normalize PLA signals to total amounts of incorporated EdU with single-cell resolution This enables to determine if a loss or gain of PLA signal is due to loss or gain of EdU proximity or of the EdU-substrate. The experimental set up of RF-SIRF thus predicts the anti-biotin EdU-PLA signal to increase with fork reversal. In contrast, the EdU-Alexa488 signal is predicted to remain unchanged regardless of fork reversal or alternatively decrease given the reported slowing of replication under situations that stall DNA replication forks ^39^ (Fig. 1a).

Low dose exposure to camptothecin (CPT, 50nM) for one hour substantially induces fork reversal and concomitantly slows replication speed, without evoking double strand breaks (DSB) as extensively validated in U2OS cells by electron microscopy (EM) and single-molecule DNA fiber assay ^1–3, 39–41^. We performed RF-SIRF under these proven fork reversal conditions (Fig. 1a). Using anti-biotin antibodies for the EdU-PLA reaction, we observed a substantial increase in EdU-PLA signals in U2OS cells challenged with CPT compared to medium-treated cells (Fig. 1b,c; median normalized EdU-PLA intensity of 0.584 with compared to 0.432 without CPT, Extended Data Fig. 1b). PLA-control reactions without EdU showed no appreciable PLA signal (Fig. 1b). In stark contrast to EdU-PLA, the EdU-Alexa488 signals decreased with CPT to some extent, consistent with the reported slowing of DNA replication upon fork reversal ^39^ (Fig. 1b,d; median normalized Alexa488 intensity of 0.21 with compared to 0.294 without CPT, Extended Data Fig. 1c). These data suggests that the increase in PLA signals seen with CPT is not caused by an inadvertent increase in EdU incorporation with CPT. Instead, the greater EdU-PLA signals evidently are caused by a closer proximity of the previously incorporated EdU moieties. To account for EdU-substrate changes with CPT treatment, we normalized the EdU-PLA signals to the EdU-Alexa488 signals on a single-cell level for an objective and robust SIRF measurement for reversed forks (RF-SIRF), which shows a significant increase in RF-SIRF signals under conditions that promote reversed forks (Fig. 1b,e; median RF-SIRF of 0.514 with CPT compared to 0.360 without, Extended Data Fig. 1d).

### Fork reversal swiftly occurs upon exposure to diverse genotoxic agents

Aside CPT, reversed replication forks are significantly induced with low doses of diverse genotoxic reagents ^22^. To test if the increase in RF-SIRF signal represents reversed forks under conditions other than CPT, we exposed U2OS cells to one hour hydroxyurea (HU, 500nM), aphidicolin (APH, 100 nM), or hydrogen peroxide (H2O2, 20 μM) as previously reported permissive for fork reversal ^22^ (Extended Data Fig. 2). When considering EdU-PLA only, H2O2 causes a significant increase in signals. Both APH and HU also show increased EdU-PLA signals, albeit without statistical significance (Extended Data Fig. 2a; median EdU-PLA intensities of 0.276 with HU, 0.292 with APH, 0.379 with H2O2 and 0.255 without external replication stress). Evidently, this was at least in part due to a loss of EdU-substrate in the treated compared to untreated cells (Extended Data Fig. 2b; median EdU-Alexa488 intensities of 0.143 with HU, 0.187 with APH, 0.227 with H2O2 and 0.419 without external replication stress). However, when calculating RF-SIRF, which takes into account the total available EdU substrate per cell, the RF-SIRF signal are significantly elevated under all RF-promoting conditions, corroborating the importance of normalization for EdU substrate availability (Extended Data Fig. 2c; median RF-SIRF of 0.342 with HU, 0.347 with APH, 0.413 with H2O2 and 0.214 without external replication stress).

Loss of nascent EdU signal amongst others is caused by loss of fork protection, which results in the degradation of nascent DNA with 1.8 kb/hour ^10^. To better control for EdU substrate changes, we tested if shorter exposures to CPT unlikely to elicit substantial degradation are sufficient to observe fork reversal by RF-SIRF. In contrast to the longer one-hour incubations with CPT, there is only a small and non-significant change in the EdU-Alexa488 signals with or without 15 minutes of CPT treatments (Figure 2a; median EdU-Alexa488 intensities of 0.293 with CPT and 0.32 without). Yet, the EdU-PLA signal intensity significantly increases under these conditions, resulting in robust RF-SIRF signals that are significantly higher when cells are challenged with CPT, thus under conditions promoting fork reversal (Fig. 2b,c; median EdU-PLA intensities of 0.432 with CPT 0.173 without). Similarly, there are smaller differences in EdU-Alexa488 signals with or without 15 minutes of HU, APH or H2O2 compared to the one-hour exposures (Fig. 2d; median EdU-488 intensities of 0.255 with HU, 0.211 with APH, 0.321 with H2O2 and 0.355 without external replication stress). In contrast to the Alexas488 intensitites, the EdU-PLA signals substantially increase with all agents tested, resulting in robust RF-SIRF signals upon exposure to the various DNA replication stresses (Fig. 2 e,f; median EdU-PLA intensities of 0.265 with HU, 0.422 with APH, 0.477 with H2O2, 0.166 without external replication stress, and median RF-SIRF of 0.252 with HU, 0.4 with APH, 0.413 with H2O2 and 0.14 without external replication stress). Taken together, the data show that RF-SIRF signals increase within short, fifteen-minute exposures to diverse genotoxic replication stalling agents known to reverse DNA replication forks, whereby oxidative damage apparently promotes the most robust RF-SIRF signals amongst the agents tested.

**Fig. 2.**
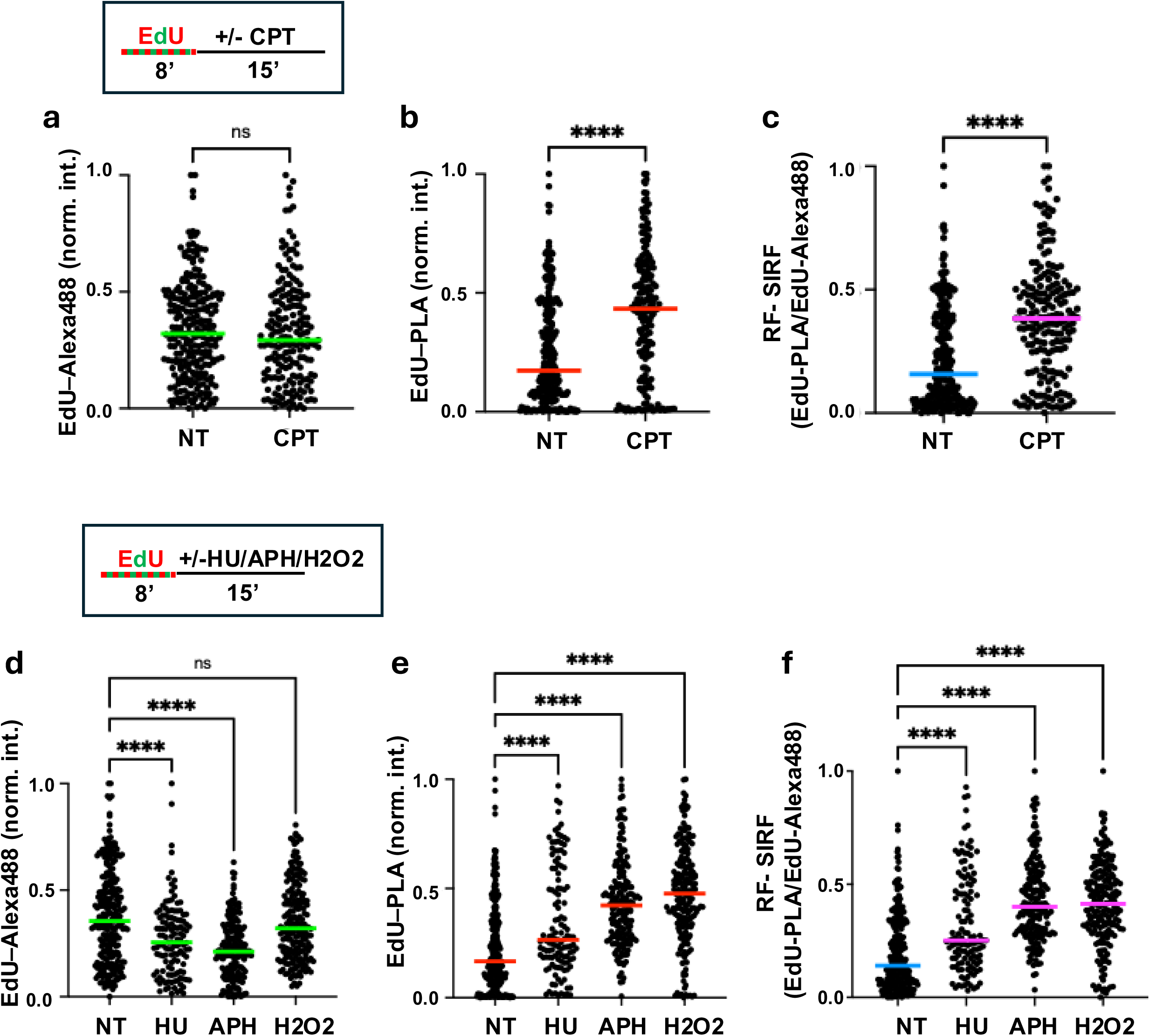
RF-SIRF signals are elevated with diverse genotoxic agents. **a.** Scatter plot of EdU-Alexa488 intensities with 15 minutes CPT (n= 189) and without CPT (NT, n= 249). Top, experimental sketch for **a-c**. **b.** Scatter plot of EdU-PLA intensities with and without with 15 minutes CPT. **c.** Scatter plot of RF-SIRF (EdU-PLA/EdU-Alexa488) with and without with 15 minutes CPT. **d.** Scatter plot of EdU-Alexa488 intensities with 15 min hydroxyurea (HU, 500μM, n= 125), aphidicolin (APH, 100nM, n= 185), hydrogenperoxide (H2O2, 20μM, n= 202) and without treatment (NT, n= 260). Top, experimental sketch for **d-f**. **e.** Scatter plot of EdU-PLA intensities with 15 min HU, APH, H2O2 and without treatment. **f.** Scatter plot of RF-SIRF (EdU-PLA/EdU-Alexa488) with 15 min HU, APH, H2O2 and without treatment. Bars denote median of each data set. Data is derived from three independent biological repeats, *p-values* are derived using the Mann-Whitney test for **a-c** and one-way ANOVA for **d-f**. \*\*\*\**p*<0.0001, \*\*\**p*<0.001, ns; not significant.

### Single-stranded (ss)DNA-SIRF signals are elevated under conditions that promote fork reversal

To further validate that RF-SIRF preferentially detects reversed replication forks, we again took advantage of the biophysical features of reversed forks. Intrinsic and unique to reversed forks, the regressed arms of reversed forks contain nascent ssDNA overhangs at various lengths ^22, 29, 40, 42^ (Figure 3a, schematic to the right, dashed box), in contrast to nascently replicated DNA at an ongoing replication fork, which is paired with its template (Figure 3a, schematic to the left). We therefore hypothesized that SIRF signals using an antibody with specificity against ssDNA ^43–45^ will increase under conditions favoring reversed forks. Consistent with this notion, ssDNA-SIRF signals significantly increased with one hour CPT treatments (Fig. 3b, median RF-SIRF of 0.476 with CPT and 0.245 without, and Extended Data Fig. 3a-d). Thus nascent ssDNA, which is limited at ongoing replication forks but known to form at reversed forks ^22, 29, 40, 42^ is elevated under conditions that promote RF-SIRF, further supporting RF-SIRF as a measure of reversed forks.

**Fig. 3.**
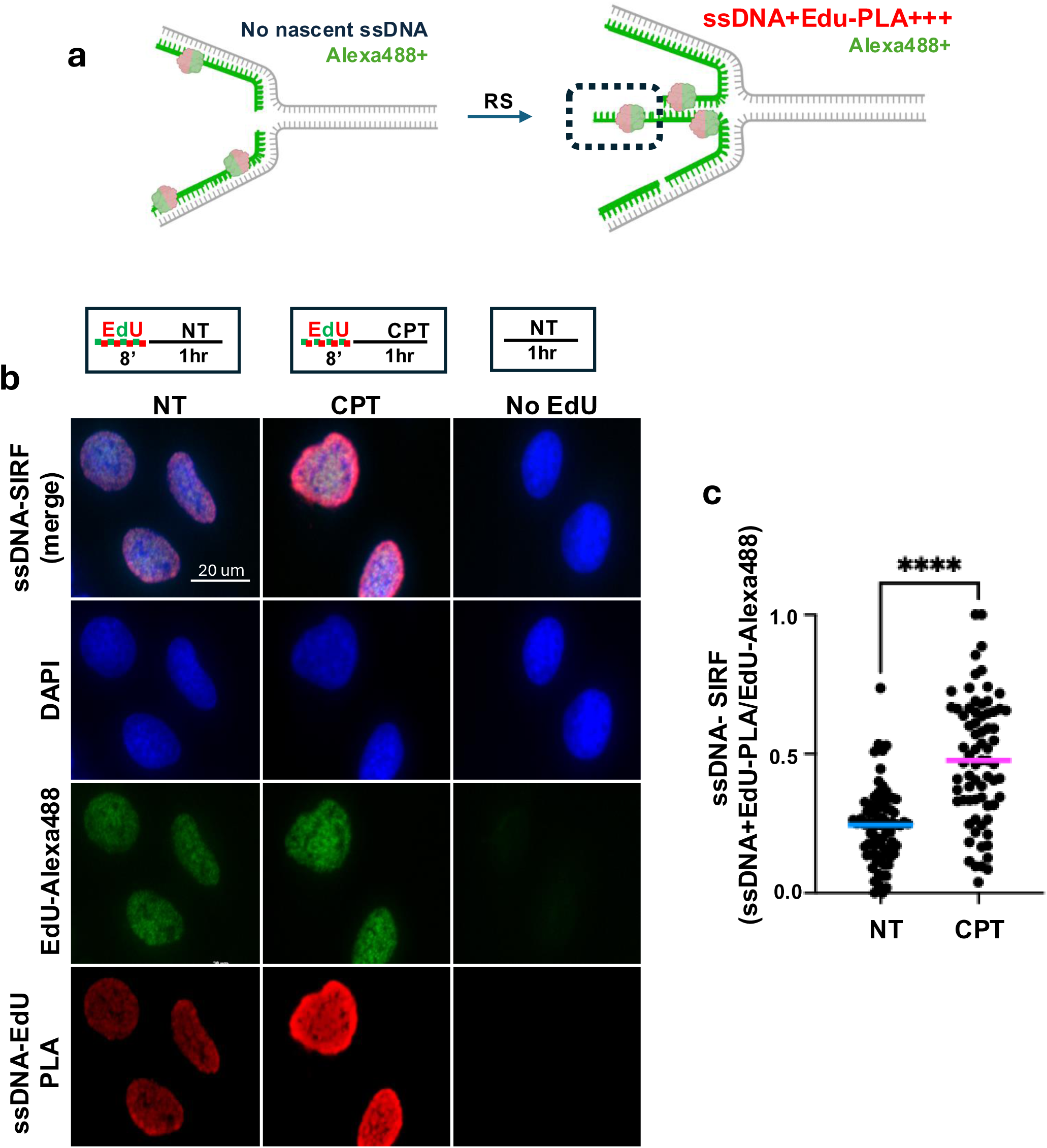
ssDNA-SIRF signals are elevated with fork reversal. **a.** Schematic of ssDNA-SIRF. Nascent single-stranded DNA (ssDNA) uniquely exists at the regressed arm of reversed DNA replication forks (right), but not at ongoing DNA replications where it is annealed to its template DNA (left). Consequently, a PLA signal using antibodies against biotinylated nascent EdU and ssDNA is expected to be increased at reversed forks compared to non-stalled forks. Sketch created with http://BioRender.com. **b.** Representative images of ssDNA-SIRF. NT (no treatment); CPT (camptothecin, 50nM, 1 hour), No EdU (negative control). Top, experimental schematic. DAPI (blue) denotes nucleus. EdU-PLA (red) shows PLA against biotinylated EdU, EdU-Alexa488 (green) shows EdU with Alexa488-Click-iT. Scale bar denotes 20μm. **c.** Scatter plot of ssDNA-SIRF (ssDNA+EdU-PLA/EdUAlexa488). NT, without replication stalling treatment (n=80), CPT, with camptothecin promoting fork reversal (n= 72). Bars denote median of each data set. Data is derived from three independent biological repeats, *p-*values are derived using the Mann-Whitney test. \*\*\*\**p*<0.0001

### RF-signals form at DNA structures controlled by SMARCAL1

SMARCAL1 is a major fork reversal DNA translocase, that dislodges nascent DNA strands at stalled DNA forks to drive the formation of reversed fork structures ^14, 20, 25^. We therefore tested the effect of SMARCAL1 ablation on RF-SIRF, which inhibits fork reversal. For this we induced reversed forks by treating U2OS cells with low-dose CPT, and compared RF-SIRF results with those in cells additionally knocked-down for SMARCAL1 (Fig. 4a, b). In direct reversal to the results obtained under conditions that promote fork reversal (Fig. 1), knocking down SMARCAL1 dramatically reduced RF-SIRF signals caused by a significant reduction in EdU-PLA signal (Fig. 4c,d; median RF-SIRF of 0.4452 with CPT and 0.2411 with CPT plus SMARCAL1 knockdown, and median EdU-PLA intensities of 0.473 with CPT and 0.315 with CPT plus SMARCAL1 knockdown). The reduction cannot be attributed to decreased EdU substrate incorporation as overall EdU-Alexa488 signals are increased, rather than decreased under SMARCAL1 knock-down conditions (Fig. 4e; median EdU-PLA intensities of 0.304 with CPT and 0.405 with CPT plus SMARCAL1 knockdown), which is consistent with accelerated DNA replication that is seen with resolution of reversed forks ^39, 46^. Similarly, HU-induced RF-SIRF signals are reduced upon SMARCAL1 knockdown, suggesting RF-SIRF signals measure reversed forks under diverse conditions (Extended Data Fig. 4a,b). Corroborating this notion, ssDNA-SIRF signals, which measure nascently replicated ssDNA characteristic of reversed forks, are reduced in cells treated with CPT or HU when SMARCAL1 expression is knocked down (Extended Data Fig. 4c-f). Taken together, the data shows that limiting reversed forks by SMARCAL1 knockdown significantly reduces RF-SIRF signals, demonstrating that RF-SIRF detects reversed DNA replication fork structures.

**Fig. 4.**
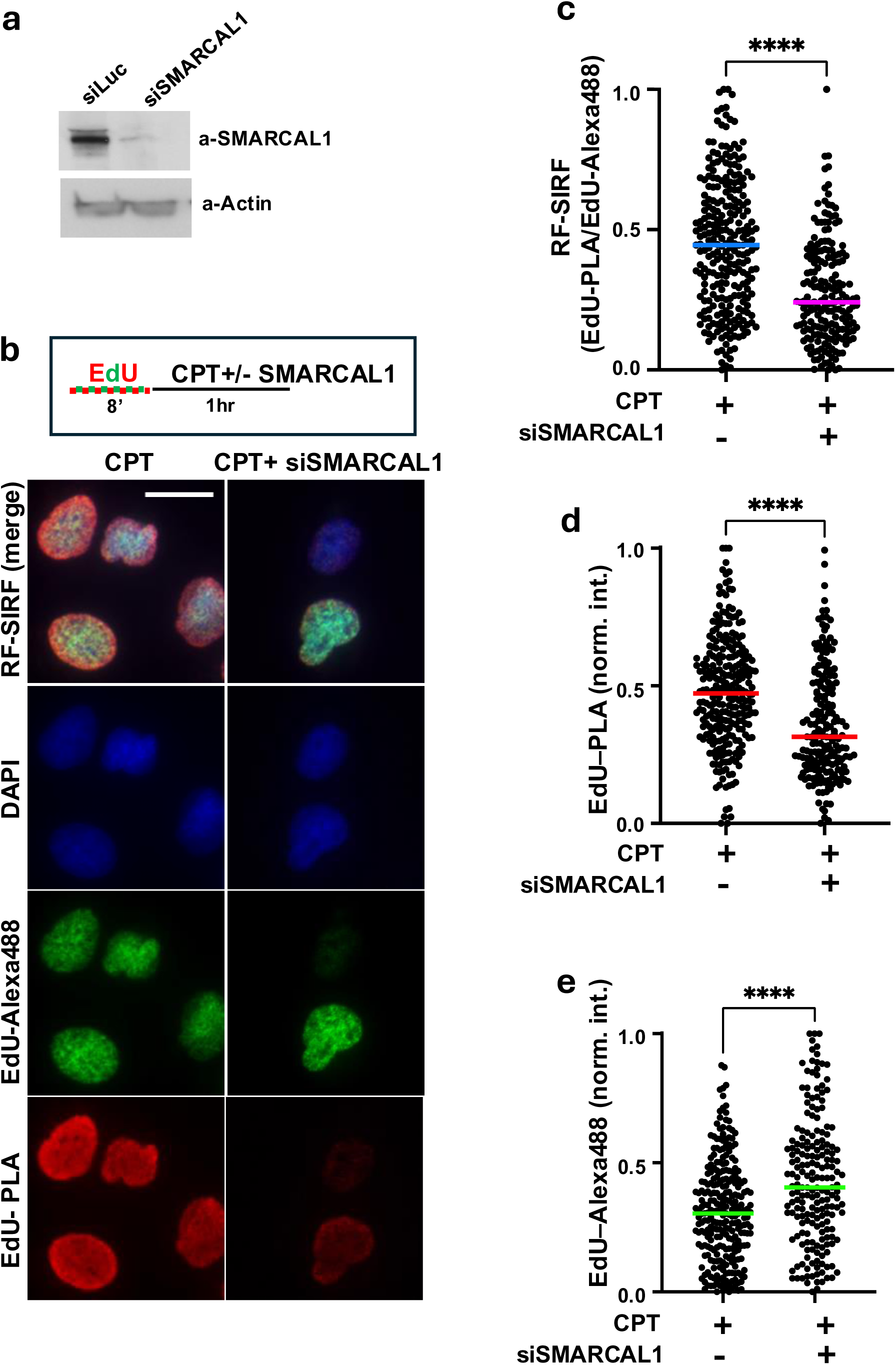
Loss of fork reversal helicase SMARCAL1 reduces RF-SIRF. **a.** Western blot of U2OS cells with SMARCAL1 knock-down (siSMARCAL1) or Luciferase knockdown as non-specific targeting control (siLuc). **b.** Representative images of RF-SIRF. CPT (camptothecin, 50nM), siSMARCAL1 (SMARCAL1 knock-down). DAPI (blue) denotes nucleus. EdU-PLA (red) shows PLA against biotinylated EdU, EdU-Alexa488 (green) shows EdU with Alexa488-Click-iT. Scale bar denotes 20μm. Top, experimental schematic. **c.** Scatter plot of EdU-PLA intensities. **d.** Scatter plot of EdU-Alexa488 intensities. **e.** Scatter plot of RF-SIRF (EdU-PLA/EdU-Alexa488). CPT, with camptothecin promoting fork reversal (n= 264), CPT+siSMARCAL1, with additional SMARCAL1 knockdown (n= 195). Bars denote median. Data is derived from three independent biological repeats, *p*-values are derived using the Mann-Whitney test. \*\*\*\**p*<0.0001

### Spatial and temporal characteristics of RF-SIRF

An advantage using a single-cell fluorescence microscopy-based approach to assess reversed forks is that it has the potential to provide temporal and spatial resolution of fork reversal reactions within cells allowing to investigate so far unexplored research questions. To further probe the RF-SIRF assay system, we performed cell-cycle analysis of RF-SIRF (Fig. 5a,b). By plotting the cell cycle profiles of cells treated with and without CPT, the data demonstrates that cells in early to mid-S-phase show the greatest increase in RF-signals in cells treated with CPT compared to untreated cells (Fig. 5a,b). Additionally, RF-signals are significantly reduced in late S-phase (Fig. 5c). Moreover, early to mid S-phase RF-signals formed with CPT or HU are reduced with SMARCAL1 knock-down (Extended Data Fig. 5a-d). Of note, while non-treated cells overall have significantly less RF-SIRF signal compared to those that are stressed by exogenously providing replication stalling agents, there are more RF signals in early-mid S-phase also in untreated cells (Extended Data Fig. 5e), suggesting that also spontaneous reversed forks occur albeit with lesser frequency in late S/G2. Together, these analyses suggest that reversed forks predominantly form in early/mid S-phase and are resolved during late-S/G2 of the cell cycle.

**Fig. 5.**
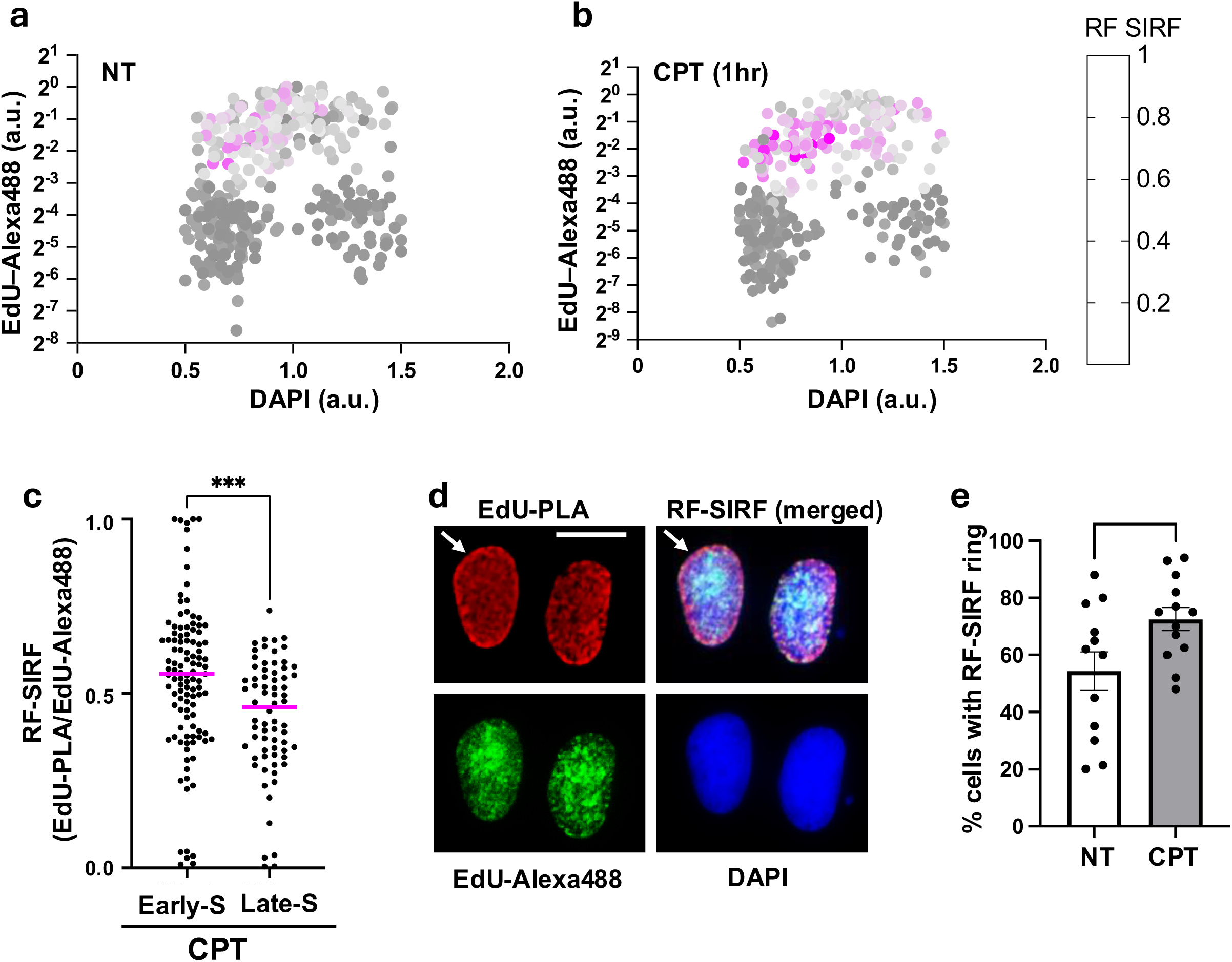
RF-SIRF signals accumulate in early/mid S-phase of the cell cycle. **a.** Multivariate scatter plots for Cell cycle analysis of RF-SIRF in untreated U2OS cells. x-axis, sum intensity of DAPI; y-axis, intensity of EdU-Alexa488 (a.u, arbitrary units, Log2 scale); the color gradient denotes RF-SIRF signals. **b.** Multivariate scatter plots for Cell cycle analysis of RF-SIRF in U2OS cells with 1 hour CPT. Data is pooled from three independent biological repeats. **c.** Scatter plot of RF-SIRF in CPT treated cells in early/mid S-phase cells (n=109), and late S-phase cells (n=70). Data pooled from three independent biological repeats, *p*-values are derived using the Mann-Whitney test. ***p<0.0001 **d.** Representative image of cells displaying EdU-PLA ring at nuclear periphery. DAPI (blue) denotes nucleus. EdU-PLA (red) shows PLA against biotinylated EdU, white arrow shows EdU-PLA ring, EdU-Alexa488 (green) shows EdU with Alexa488-Click-it. Scale bar denotes 20μm. **e.** Bar graph of percentage of cells per image containing EdU-PLA ring at nuclear periphery with and without CPT (50nM, 1 hour). Bars denote mean, error bars the standard error of the mean. Data is derived from three independent biological repeats, *p*-values are derived using the two-tailed unpaired t-test. \**p*<0.01

Stalled DNA replication forks can be tethered to nuclear pores at the nuclear periphery ^47–49^. In addition to temporal resolution, we sought to query the subcellular localization of RF-signals to obtain spatial knowledge of reversed forks. We noticed that RF-SIRF often forms a prominent signal at the nuclear periphery in proximity to the nuclear lamina, which is in contrast to EdU-Alexa488 signals (Fig. 5d, Extended Data Fig. 5f). Moreover, the number of cells containing prominent peripheral nuclear RF-ring like structures increase significantly with treatment of CPT (Fig. 5e, 72.6% cells with nuclear ring-like RF-signal with CPT compared to 54.4% without). These data suggest that reversed forks preferentially albeit not exclusively form at the nuclear periphery.

### Local chromatin changes at reversed replication forks

While canonically considered for transcriptional control, local chromatin changes during DNA stress are critical intracellular communication markers enabling the specific recruitment of DNA stability factors ^17, 34, 35, 49–51^. Whether fork reversal reprograms the epigenetic landscape is currently elusive. Having established a robust assay system to interrogate reversed fork reactions, we sought to test its usage for understanding potential epigenetic changes that may occur at reversed forks (Fig. 6a). RF-SIRF signals show a preference for the nuclear periphery and are more prevalent during early/mid S-phase of cell cycle, a pattern shared with heterochromatin ^49, 52, 53^. H3K9me3 is a heterochromatin mark known to be induced with replication stress. To test if heterochromatin forms at reversed forks, we performed H3K9me3-SIRF with and without CPT (Fig. 6b,c, and Extended Data Fig. 6a,b). Under conditions promoting fork reversal, H3K9me3-SIRF signals are significantly enriched (Fig. 6c, median H3K9me3-SIRF of 0.347 with CPT and 0.091 without CPT). Importantly, H3K9me3-SIRF signals are reduced to non-stressed levels when knocking down SMARCAL1 (Fig. 6c, median H3K9me3-SIRF of 0.347with CPT and 0.049 without CPT plus SMARCAL1 knockdown). These data demonstrate that H3K9me3 is enriched at reversed replication forks, and not at nascent DNA behind the fork, which is not expected to change with SMARCAL1 knock down.

**Fig. 6.**
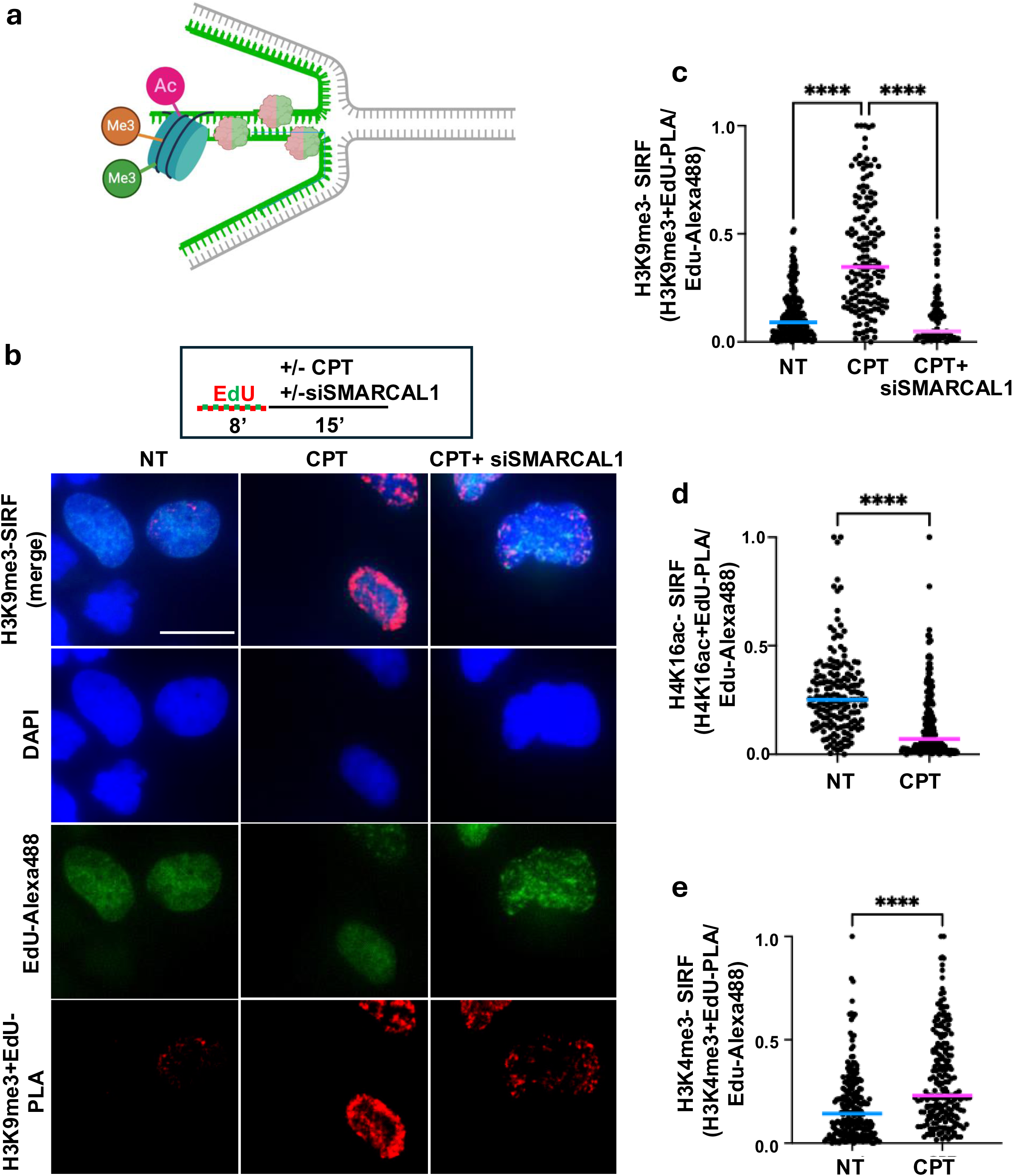
Local chromatin at reversed forks. **a.** Schematic of chromatin-SIRF. SIRF using antibodies against epigenetic markers of interest under conditions that reversed forks allows testing chromatin at reversed forks. Sketch created with http://BioRender.com. **b.** Representative images of H3K9me3-SIRF. NT (no treatment); CPT (camptothecin, 50nM, 15 minutes), CPT+siSMARCAL1 (with SMARCAL1 knockdown as negative control for reversed forks). Top, experimental schematic. DAPI (blue) denotes nucleus. H3K9me3-PLA (red) shows PLA against H3K9me3 and biotinylated EdU, EdU-Alexa488 (green) shows EdU with Alexa488-Click-iT. Scale bar denotes 20μm. **c.** Scatter plot of H3K9me3-SIRF (H3K9me3+EdU-PLA/EdUAlexa488). NT, without replication stalling treatment (n=202), CPT, with camptothecin promoting fork reversal (n= 154). CPT+siSMARCAL1 (with knockdown of SMARCAL1, n= 75). Data pooled from 2-4 independent biological repeats. **d.** Scatter plot of H4K16ac-SIRF (H4K16ac+EdU-PLA/EdUAlexa488). NT, without replication stalling treatment (n=174), CPT, with camptothecin promoting fork reversal (n= 171). Data pooled from 2 independent biological repeats. **e.** Scatter plot of H3K4me3-SIRF (H3K4me3+EdU-PLA/EdUAlexa488). NT, without replication stalling treatment (n=232), CPT, with camptothecin promoting fork reversal (n= 214). Data pooled from 3 independent biological repeats. Bars denote median of each data set. Data is derived from three independent biological repeats, *p*-values are derived using the Mann-Whitney test. \*\*\*\**p*<0.0001

H4K16ac is an epigenetic mark for transcriptionally active chromatin. In the context of DNA stress responses, it is a critical mark for suppressing TP53BP1 binding to damaged DNA sites ^51, 54^. TP53BP1 is also recruited to stalled replication forks and acts in replication fork protection ^21^. Under the same fork reversal promoting conditions that show an increase in H3K9me3, H4K16ac-SIRF signals are significantly decreased (Fig. 6d, median H4K16ac-SIRF of 0.070 with CPT and 0.251 without CPT, and Extended Data Fig. 6c,d). These data show that distinct epigenetic chromatin changes occur under conditions that promote fork reversal.

Similar to H4K16ac, H3K4m3 is an epigenetic mark for transcriptionally active chromatin. Yet in the context of DNA stress responses, it is required for MRE11 recruitment to stalled replication forks suppressing cancer therapy resistance in BRCA1/2 defective cancer cells ^17, 35^. Intriguingly, despite both H4K16ac and H3K4me3 marking active transcription, they show opposite behavior at reversed replication forks with CPT, whereby H3K4me3-SIRF signals are significantly enriched at CPT-stalled forks (Fig. 6e, median H3K4me3-SIRF of 0.23 with CPT and 0.143 without CPT, and Extended Data Fig. 6e,f). Consistent with previous data implying MRE11 to act at reversed fork ^10–15^, these data explain how MRE11 is recruited there. Taken together the data shows that the epigenetic code for replication stress differs from that at transcription sites and explains previously observed proteomic changes at reversed forks.

## Discussion

Here we describe RF-SIRF, an in-situ method for detecting reversed replication forks with single-cell resolution. The assay exploits the unique DNA structures formed at reversed forks and leverages from distinct experimental conditions extensively verified to promote fork reversal. It is a robust, efficient and reproducible technique that can be performed with minimal processing steps using standard molecular and microscopy techniques. As a productive EdU-PLA signal depends on both, the proximity and the total amount of EdU present in the DNA, it is important to note that proper, context-dependent controls should be included to accurately interpret RF-EdU. This is particularly of importance when large changes in EdU-Alexa488 are observed between conditions, which could be the result of fork degradation or fork slowing. Nevertheless, our data suggest that conditions with short-time, acute replication stalling induces robust fork reversal while largely circumventing substantial EdU changes and therefore provides a condition for added confidence in assessing reversed forks.

Our data demonstrate that RF-SIRF constitutes a new tool that provides spatial context with regards to subcellular localization of reversed forks as well as proteomic changes associated with reversed forks. It has been shown that stalled DNA replication forks relocate to nuclear pores at the nuclear periphery ^47^. RF-SIRF corroborates this notion by demonstrating that reversed replication forks preferentially form at the nuclear periphery, a location shared with canonical heterochromatin and lamins. Consistent with a particular role for reversed forks at the periphery, the progeroid disease suppressor gene *LMNA*, encoding for Lamin A/C proteins, has been reported to protect stalled forks from degradation ^55^, a process that primarily occurs at reversed replication forks ^10, 13 12, 14–16^. Nevertheless, it is to note that RF-signals also form inside the lumen of the nucleus. It will be interesting to explore genome-wide RFs to study co-localization with centromeric regions, telomers and common fragile sites, enabled by the in-situ nature of RF-SIRF.

So far it remained elusive if chromatin forms at a reversed fork, which given the importance of epigenetic modifications in the recruitment of DNA stress response proteins to damaged DNA sites, has been of great interest. Somewhat paradoxical to this, it was reported that heterochromatin assembles at nascent DNA ^52^, which canonically is associated with chromatin compaction that makes protein access unamicable. Yet MRE11 acts at reversed forks ^10, 12–14^. Here, the SIRF data here demonstrating that H3K4me accumulates at reversed forks explains how MRE11 is recruited there and is consistent with its biophysical property as a DNA end processing protein. Notably, the data shows that both transcriptionally active and repressive marks are induced at reversed forks, suggesting that these marks may not necessarily reflect chromatin compaction states at stalled forks. Instead, it suggests that a distinct epigenetic language exists at stalled replication forks, that differs from the coding canonically utilized during transcription to mark active and repressed states. It will be important to better understand the epigenetic replication code in future studies.

Collectively, RF-SIRF enables a valuable robust and effective opportunity to study reversed replication forks within their cellular, spatial and proteomic context in cells to inform on molecular mechanisms of DNA stress responses critical during cancer therapy responses, development and disease suppression.

## Methods

### Reagents, cell lines and culture conditions

Hydroxyurea (HU), Aphidicolin, Hydrogen Peroxide, X-tremeGENE^™^ HP DNA Transfection Reagent and Duolink proximity ligation assay reagents were from Sigma-Aldrich; Ethynyl-2’-deoxyuridine (EdU), biotin azide, Alexa Fluor 488 azide, Alexa Fluor 647 azide and ProLong Gold were from Invitrogen; 4′,6-diamidino-2-phenylindole (DAPI, 62248) and Lipofectamine™ RNAiMAX Transfection Reagent was obtained from ThermoFisher Scientific; 32% Paraformaldehyde (PFA) was from Electron Microscopy Science, protein block buffer was from Abcam, Camptothecin was from Selleck chemicals, T4 DNA ligase buffer from New England Biolabs. Antibodies used for SIRF assays are as follows: mouse anti-biotin (Sigma-Aldrich), rabbit anti-biotin (Cell Signaling Technology), mouse anti-dsDNA (Millipore), mouse anti-dsDNA (Millipore), rabbit anti-GFP (Cell signaling technology), rabbit anti-H3K9me3 (Abcam), mouse anti-ssDNA (Millipore), mouse anti-Lamin A/C (Cell signaling technology) rabbit anti-H3K4me3 (Abcam), rabbit anti-H4K16ac (Abcam). Antibodies used for immunoblotting are mouse anti-SMARCAL1 (Santa Cruz Biotechnology) and mouse anti-beta-actin (sigma). SMARCAL1 siRNA was obtained from Horizon. U2OS cells (ATCC) were grown in Dulbecco’s Modified Eagle Medium (DMEM, Life Technologies) supplemented with 10% fetal bovine serum (Gemini Bio-Products).

### Western blot for siRNA

SMARCAL1 siRNA transfection was performed using RNAiMAX following manufacturer’s instructions. Cell lysates were resolved by SDS-PAGE. Proteins were transferred to PVDF membranes, and incubated with primary antibodies overnight. Corresponding secondary antibodies were incubated for 1 hr at RT. Signals were detected using enhanced chemiluminescence.

### RF-SIRF protocol

Cells are grown in log-phase and plated the day before the experiment onto microscope chamber slides. Cells are treated with EdU, washed with Phosphate buffered saline (PBS, pH7.4), followed by addition of media (no treatment), or genotoxic agent for 1 hour or 15 minutes, as indicated in the figures. Cells are fixed, permeabilized and a Click-iT reaction is performed according to manufacturer’s instructions. After blocking and primary antibody incubation, Duolink PLA (Sigma) is performed according to manufacturer’s instructions with an additional incubation step before incubation with DAPI and mounting with Prolong Gold antifade (Life Technologies). Slides are imaged using a Nikon Eclipse Ti-U inverted microscope and analyzed using Nikon NIS elements software.

### Image acquisition and analyses

Images are obtained using Nikon Eclipse Ti-U inverted microscope with Andor Zyla camera and Plan Apochromat objective lens at 40X magnification. 3-6 image fields were acquired for each condition, yielding 100-300 nuclei for image analyses per condition and biological independent experiment. Mean fluorescent intensity for DAPI, GFP, Texas-red and Cy5 channels is determined using NIS elements software. RF-SIRF and SIRF values are calculated using the ratio between PLA intensity and corresponding EdU-Alexa488 intensity per cell. Fluorescent intensity data is min-max normalized for each independent biological experiment. The Mann-Whitney test is applied to determine statistical significance when two variables are being compared. The one-way ANOVA is applied to determine statistical significance when more than two variables are being compared.

## Data availability

All data pertaining to the results in the manuscript are available in the main text, the Supplementary materials, and Source Data file. Any additional data and material requests are required to comply with institutional policies and can be requested by contacting the corresponding authors upon peer reviewed publication.

## Acknowledgements

We thank Mr. Robert Darling and Ms. Katerina Kourpas for efforts on RF experiments before final development of the current assay. The work was supported by the NIEHS under award 1R01ES029680, and by CPRIT RP180463, R1312 and RP180813 (K.S.), bridge funds by the Department of Cancer Biology UT MD Anderson Cancer Center. Morgan Fimreite was supported by the CPRIT Research Training Award CPRIT Training Program (RP210028). K.S. is a Rita Allen Foundation Fellow and a CPRIT scholar in Cancer Biology (previous award R1312).

## Notes

### Competing Interest Statement

The authors have declared no competing interest.

