## supplemental figures and legend for "RF-SIRF defining reversed DNA replication forks with single-cell and spatio-temporal resolution reveals a replication stress specific epigenetic code"

Extended Data Figures

#### Extended Data Fig. 1

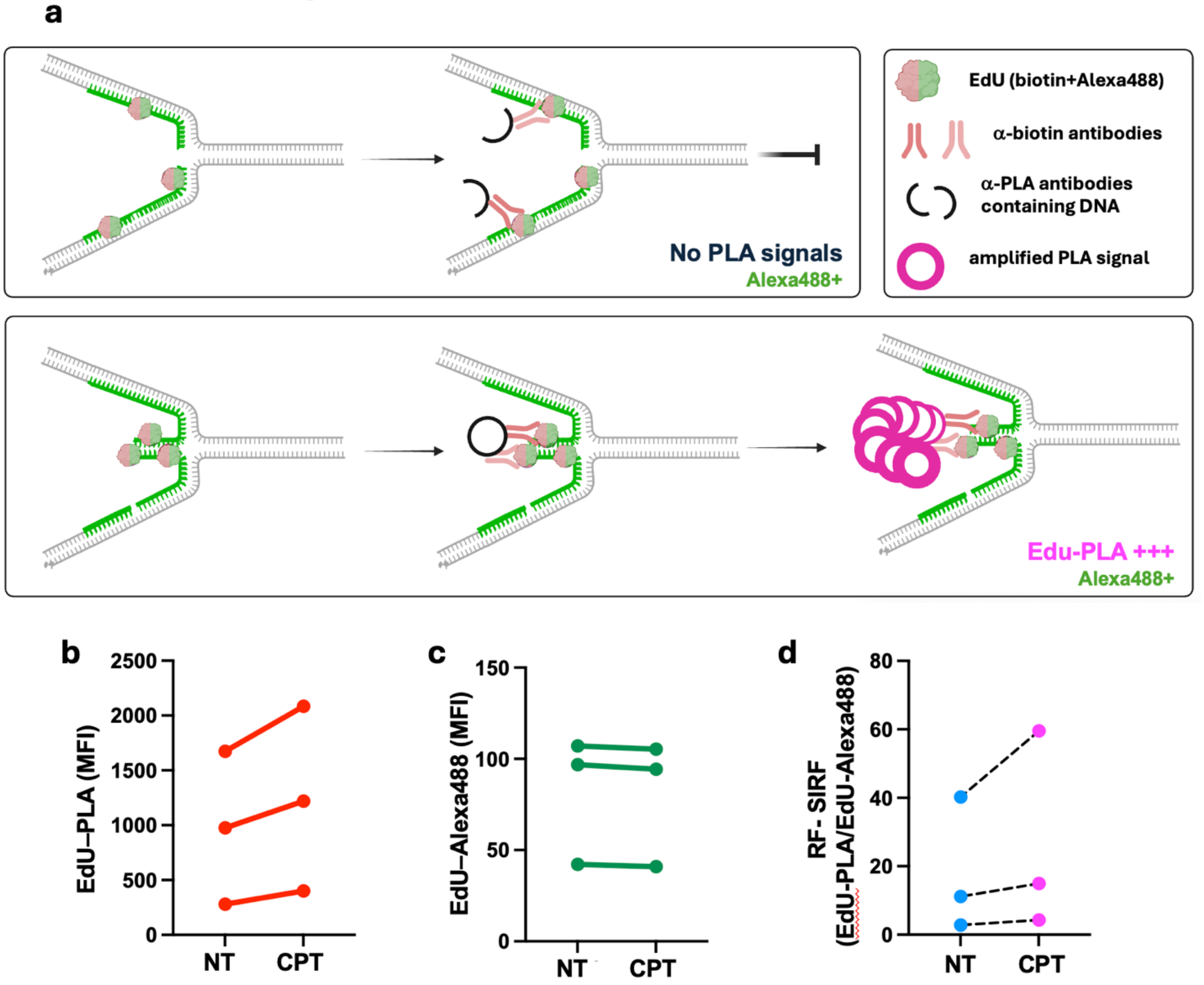


**Extended Data Fig. 1. RF-SIRF measures proximity of EdU at reversed forks**
**a.** Detailed schematic of RF-SIRF. Nascently incorporated EdU residues are simultaneously labeled with biotin and Alex488 using Click-iT chemistry (red and green bubbles). The intensity of the Click-iT reaction with Alexa Fluor 488-azide is proportional to the amount of EdU substrate, hence directly measuring the total incorporated EdU. In contrast, the proximity ligation assay (PLA) assesses the spatial proximity between two epitopes of interest. In RF-SIRF, biotinylated-EdU is recognized by anti-biotin primary antibodies (red antibodies). PLA secondary antibodies contain a DNA oligomer (black half circle) that when in close proximity to each other (<40nm), can be ligated to form a DNA circle (black circle). Subsequent rolling circle replication of the circular DNA allows for a ~100fold amplification of DNA. By annealing a fluorescent DNA probes, the resultant fluorescent signal is thus amplified so that a single protein-protein interaction produces a robust fluorescent signal detectable by standard fluorescence microscopy (pink circles). Notably, the production of the fluorescent signal depends on the proximity

of the primary epitopes. Without rolling circle amplification of a DNA circle, which directly depends on the proximity of the primary epitopes, no PLA signal is produced. Thus, at reversed forks, where the nascently incorporated EdU moieties are in close proximity within the regressed arm of the annealed nascent leading and lagging strand (green DNA) and are expected to produce robust PLA signal (bottom sketch) less likely to occur at a canonical Y-fork replication fork (top sketch). Importantly, EdU-PLA signals are compared and normalized to the total amount of incorporated EdU as measured by EdU-Alexa488 to differentiate between a signal changes caused by changes in PLA substrates (e.g. amounts of nascent DNA and EdU substrate) from a loss of actual protein-DNA interaction.
**b.** Mean of absolute EdU-PLA intensities for three biologically independent experiments with CPT and without (NT). Connecting line for visualization of median change.
**c.** Mean of absolute Alexa488 intensities for three biologically independent experiments with CPT and without (NT). Connecting line for visualization of median change.
**d.** Mean of EdU-PLA-Alexa488 ratios using absolute intensities for three biologically independent experiments with CPT and without (NT). Connecting line for visualization of median change.

#### Extended Data Fig. 2

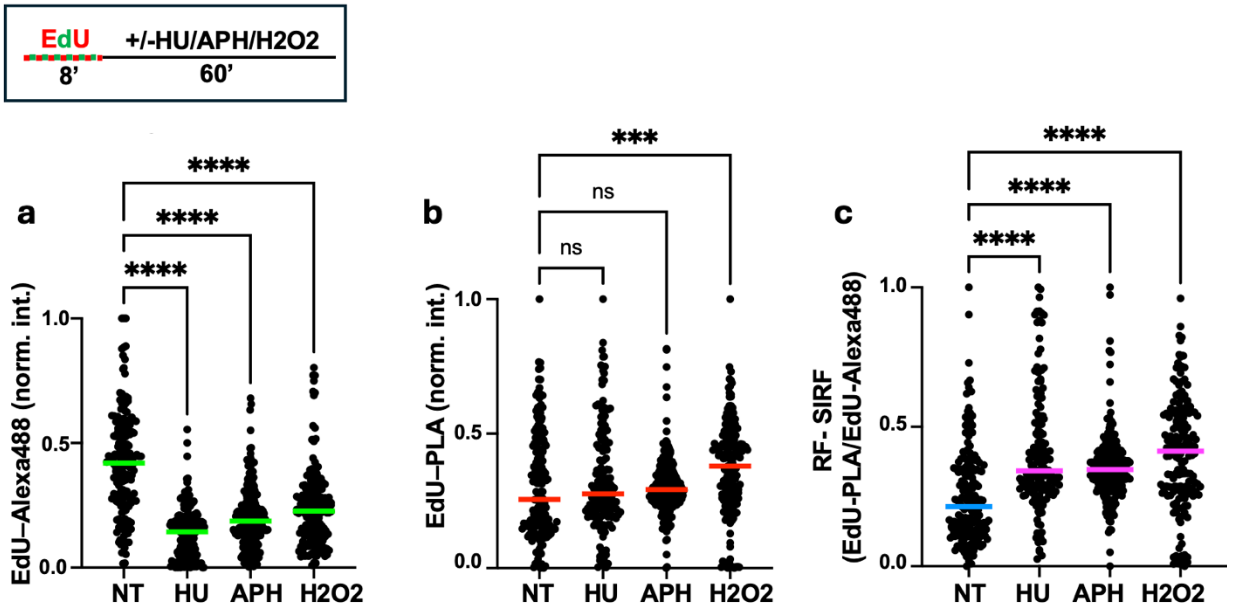


**Extended Data Fig. 2 RF-SIRF is elevated under diverse replication stalling conditions known to cause fork reversal**

**a.** Scatter plot of EdU-PLA intensities with 1 hour HU (500μM, n= 161), APH (100nM, n= 208), H2O2 (20μM, n= 172) and without treatment (NT, n= 177). Top, experimental sketch for **a-c**.

**b.** Scatter plot of EdU- Alexa488 intensities with 1 hour HU, APH, H2O2 and without treatment.

**c.** Scatter plot of RF-SIRF (EdU-PLA/EdU-Alexa488) with 1 hour HU, APH, H2O2 and without treatment.

Bars denote median of each data set. Data is derived from three independent biological repeats, *p*-values are derived using the one-way ANOVA. ****$p<0.0001$, ***$p<0.001$, ns; not significant.

#### Extended Data Fig. 3

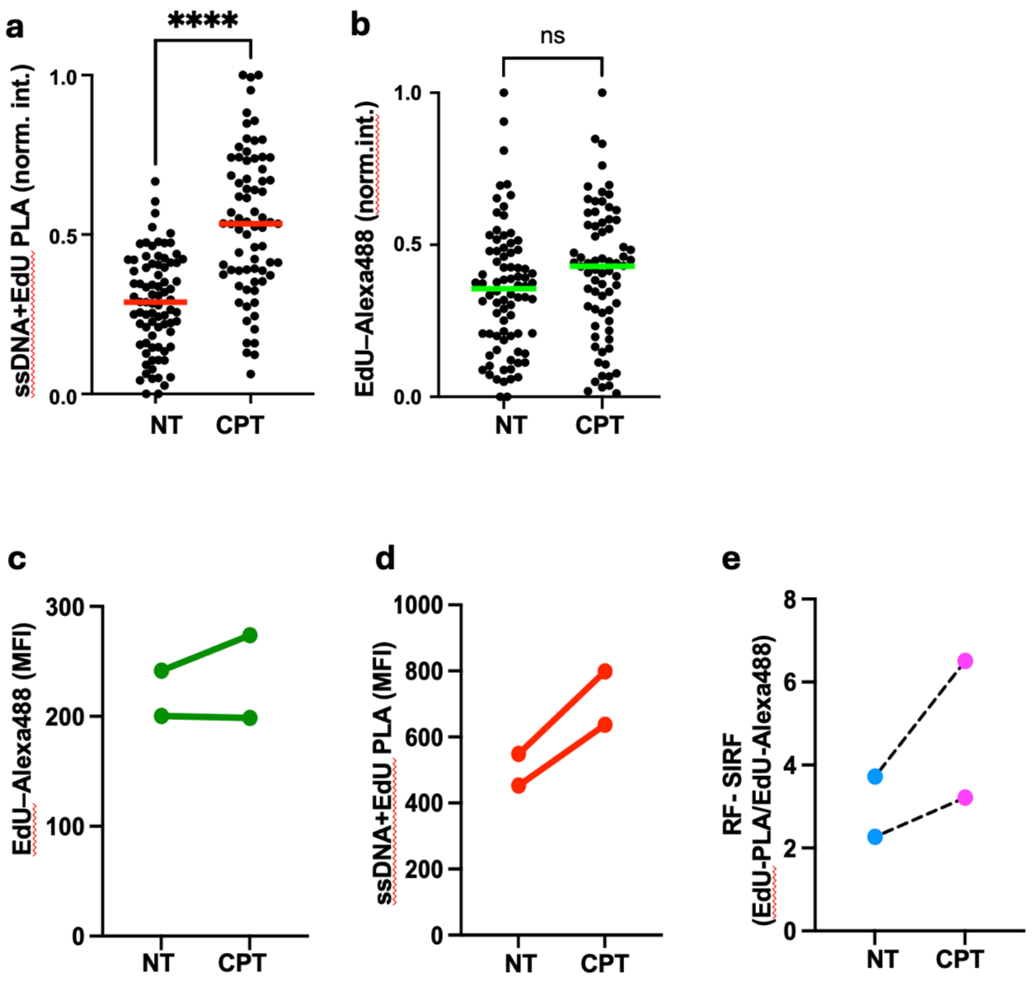


**Extended Data Fig. 3 ssDNA+EdU-PLA are elevated with fork reversal**
**a.** Scatter plot of ssDNA+EdU-PLA intensities with and without 1 hour CPT.
**b.** Scatter plot of EdU-Alexa488 intensities.
**c.** Mean of absolute ssDNA-PLA intensities for two biologically independent experiments with CPT and without (NT). Connecting line for visualization of median change.
**d.** Mean of absolute Alexa488 intensities for two biologically independent experiments with CPT and without (NT). Connecting line for visualization of median change.
**e.** Mean of ssDNA+EdU-PLA : Alexa488 ratios using absolute intensities for two biologically independent experiments with CPT and without (NT). Connecting line for visualization of median change.
NT, without replication stalling treatment (n=80), CPT, with camptothecin promoting fork reversal (n= 72). Bars denote median of each data set. Data is derived from three independent biological repeats, *p*-values are derived using the Mann-Whitney test. ****$p<0.0001$, ns; not significant.

### Extended Data Fig. 4

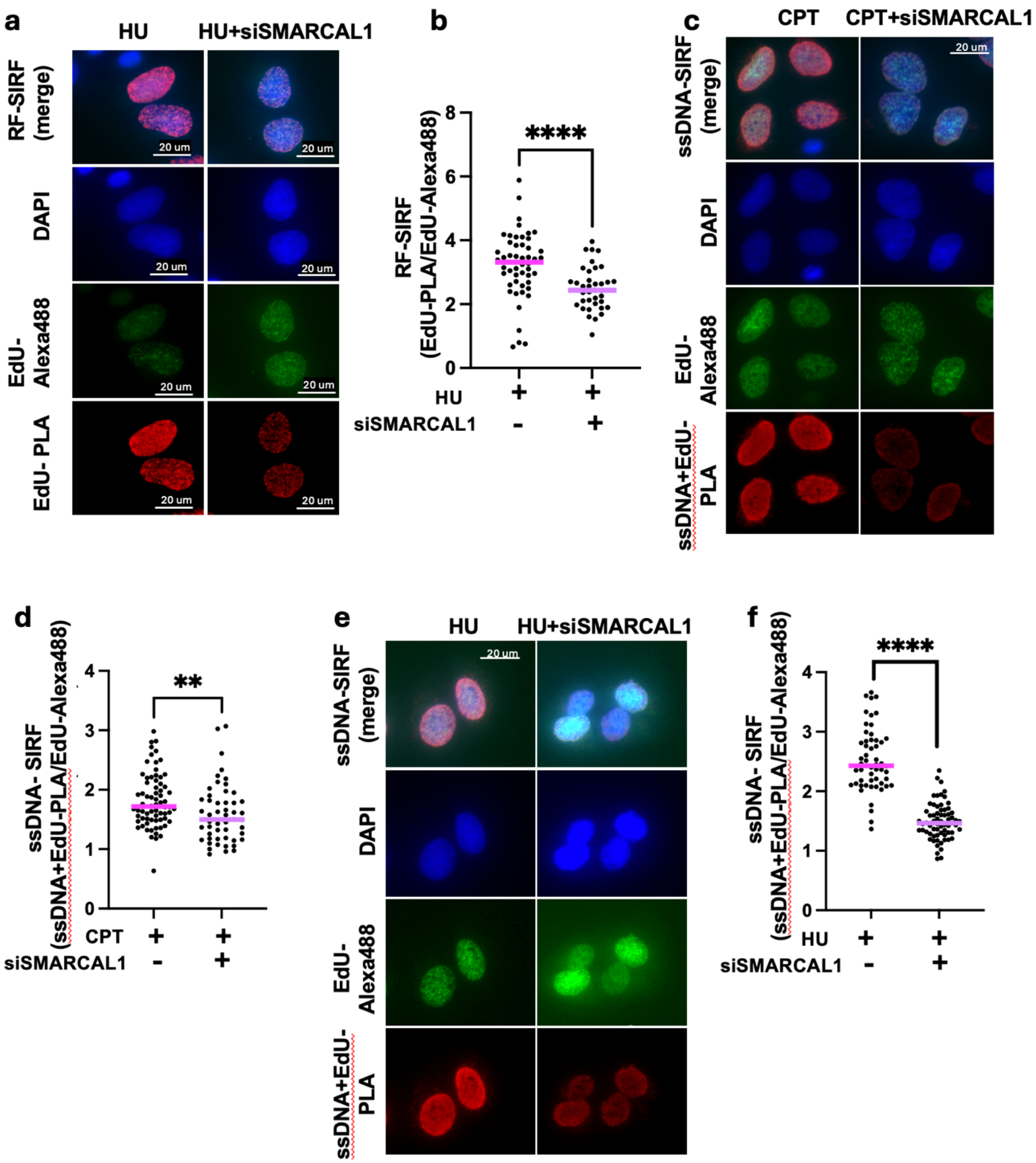


**Extended Data Fig. 4. Loss of fork reversal helicase SMARCAL1 reduces ssDNA-SIRF**

**a.** Representative images of RF-SIRF with hydroxyurea (HU, 500μM 1hour), with and without siSMARCAL1 (SMARCAL1 knock-down). DAPI (blue) denotes nucleus. EdU-

PLA (red) shows PLA against biotinylated EdU, EdU-Alexa488 (green) shows EdU with Alexa488-Click-it. Scale bar denotes 20μm.

**b.** Scatter plot of RF-SIRF (EdU-PLA/EdU-Alexa488). HU, with hydroxyurea promoting fork reversal (n= 51), HU+siSMARCAL1, with additional SMARCAL1 knockdown (n= 37).

**c.** Representative images of ssDNA-SIRF with camptothecin (CPT, 50nM, 1 hour), with and without siSMARCAL1 (SMARCAL1 knock-down). DAPI (blue) denotes nucleus. ssDNA-PLA (red) shows PLA against ssDNA and biotinylated EdU, EdU-Alexa488 (green) shows EdU with Alexa488-Click-it. Scale bar denotes 20μm.

**d.** Scatter plot of ssDNA-SIRF (EdU-PLA/EdU-Alexa488). CPT (n= 70), CPT+siSMARCAL1 (n= 51).

**e.** Representative images of ssDNA-SIRF with hydroxyurea (HU, 500μM, 1 hour), with and without siSMARCAL1 (SMARCAL1 knock-down). DAPI (blue) denotes nucleus. ssDNA-PLA (red) shows PLA against ssDNA and biotinylated EdU, EdU-Alexa488 (green) shows EdU with Alexa488-Click-it. Scale bar denotes 20μm.

**f.** Scatter plot of ssDNA-SIRF (EdU-PLA/EdU-Alexa488). HU (n= 55), HU+siSMARCAL1 (n=67).

Bars denote median. Data is shown from one biological repeat. *p*-values are derived using the Mann-Whitney test. ****$p<0.0001$, **$p<0.01$

#### Extended Data Fig. 5

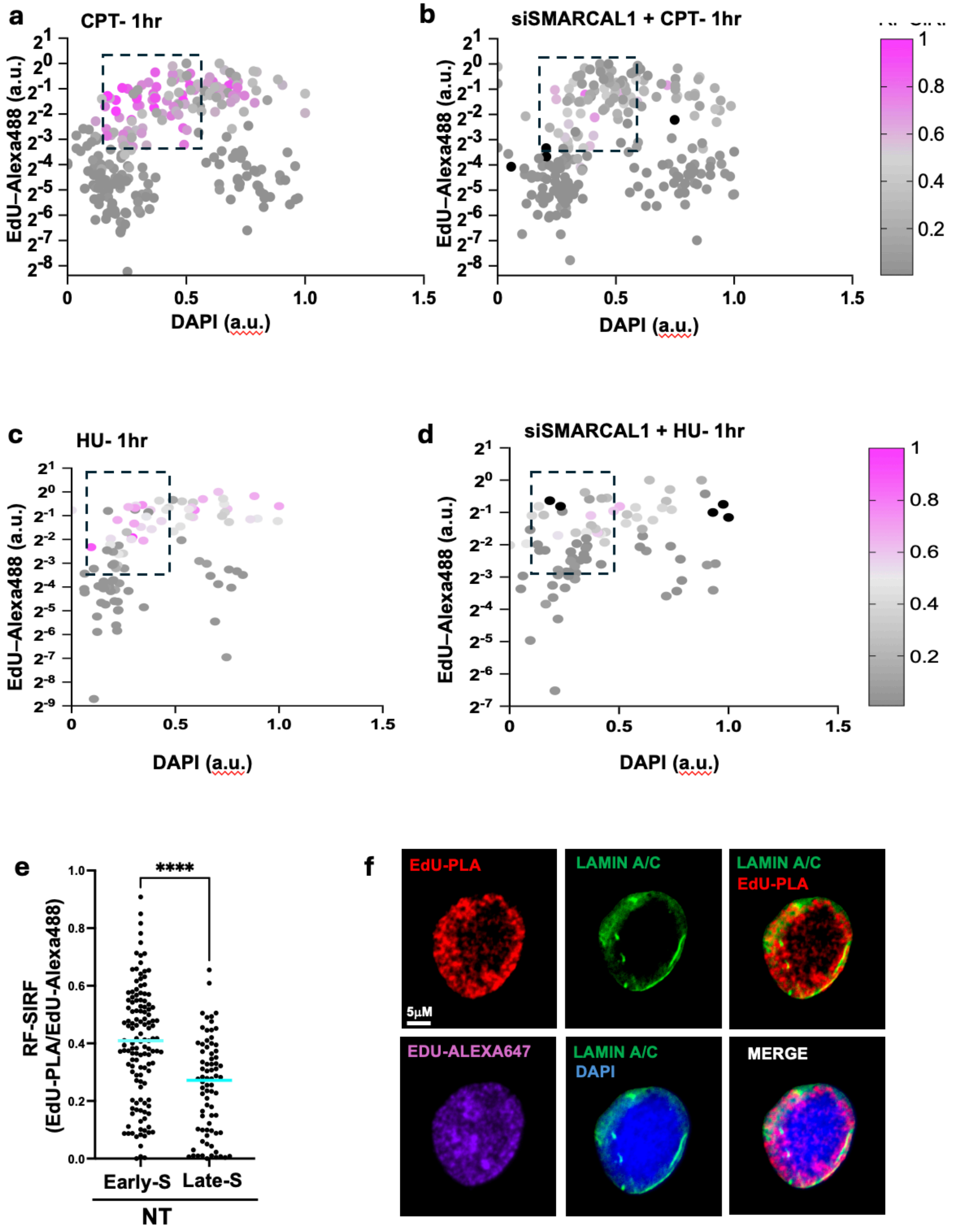


**Extended Data Fig. 5. RF-SIRF signals accumulate in early/mid S-phase of the cell cycle**

**a.** Multivariate scatter plots for Cell cycle analysis of RF-SIRF in CPT treated U2OS cells. x-axis, sum intensity of DAPI; y-axis, intensity of EdU-Alexa488 (Log2 scale); the color gradient denotes RF-SIRF signals.
**b.** Multivariate scatter plots for Cell cycle analysis of RF-SIRF in CPT treated U2OS with SMARCAL1 knock down (siSMARCAL1). Data is pooled from two independent biological repeats.
**c., d.** Multivariate scatter plots for Cell cycle analysis of RF-SIRF in hydroxyurea (HU) treated U2OS with SMARCAL1 knock down (siSMARCAL1, (**d**)) or without (**c**).
**e.** Scatter plot of RF-SIRF in non-treated cells (NT) in early/mid S-phase cells (n=129), and late S-phase cells (n=76). Data pooled from three independent biological repeats, *p*-values are derived using the Mann-Whitney test. ***$p<0.0001$
**f.** Representative image of cells displaying EdU-PLA ring at nuclear periphery co-stained with LaminA. DAPI (blue) denotes nucleus. EdU-PLA (red) shows PLA against biotinylated EdU, EdU-Alexa647 (magenta) shows EdU with Alexa647-Click-iT, green shows LaminA immunofluorescence. Scale bar denotes 5μm.

#### Extended Data Fig. 6

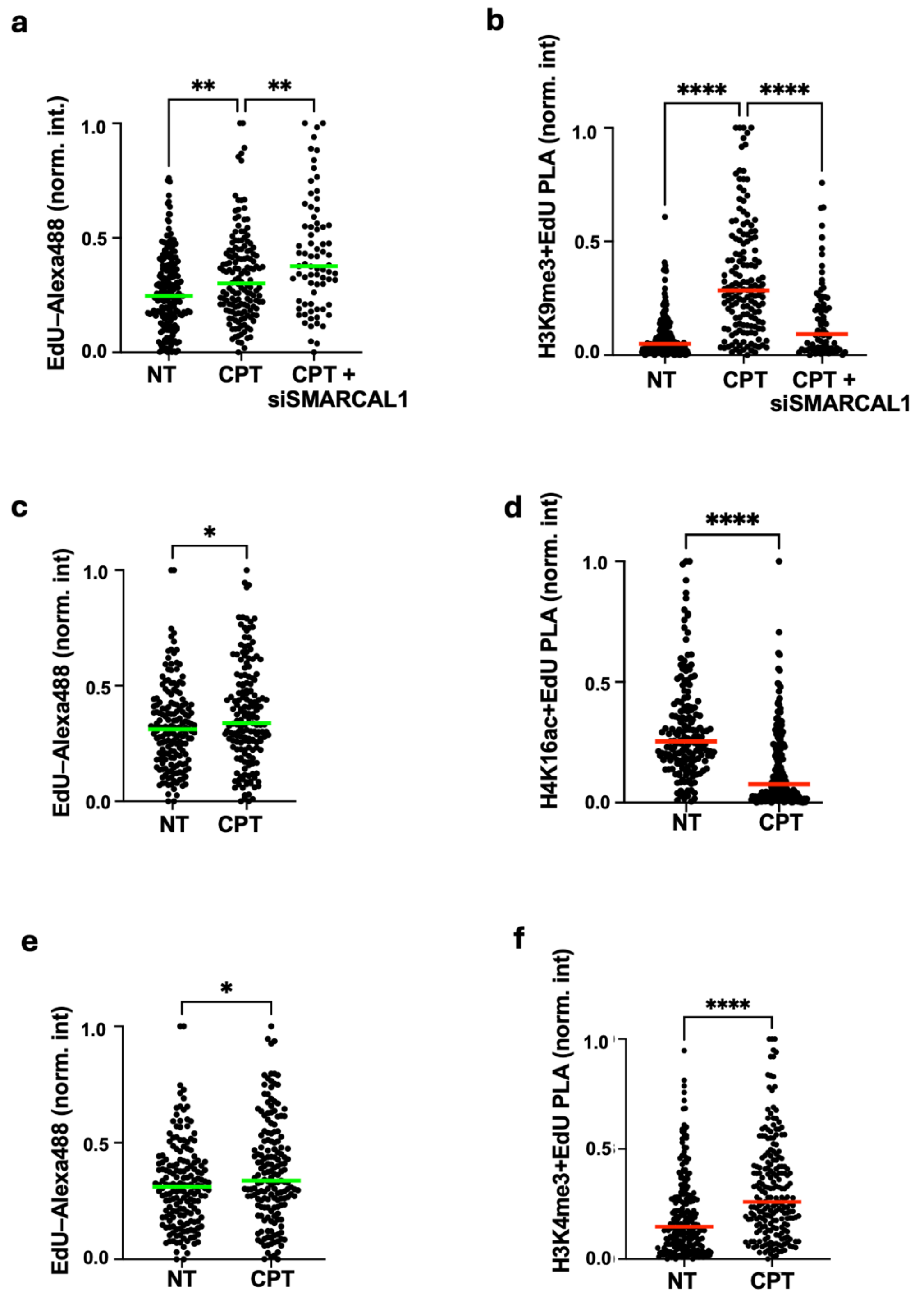


**Extended Data Fig. 6** Local chromatin-PLA is changed with camptothecin

**a,b.** Scatter plot of H3K9me3-PLA (**a**) and EdU-Alexa488 (**b**). NT, without replication stalling treatment (n=202), CPT, with camptothecin promoting fork reversal (n= 154). CPT+siSMARCAL1 (with knockdown of SMARCAL1, n= 75).
**c,d.** Scatter plot of H4K16ac-PLA (**c**) and EdU-Alexa488 (**d**). NT, without replication stalling treatment (n=174 CPT, with camptothecin promoting fork reversal (n= 171).
**e,f.** Scatter plot of H3K4me3-PLA (**e**) and EdU-Alexa488 (**f**). NT, without replication stalling treatment (n=232), CPT, with camptothecin promoting fork reversal (n= 214)
Bars denote median of each data set. Data is derived from three independent biological repeats, *p*-values for **a,b** are derived using a one-way ANOVA, and for **c-f** using the Mann-Whitney test. ****$p<0.0001$, **$p<0.01$, ns, not significant.
